# Prediction of fluoroquinolone susceptibility directly from whole genome sequence data using liquid chromatography-tandem mass spectrometry to identify mutant genotypes

**DOI:** 10.1101/138248

**Authors:** Wan Ahmad Kamil Wan Nur Ismah, Yuiko Takebayashi, Jacqueline Findlay, Kate J. Heesom, Juan-Carlos Jiménez-Castellanos, Jay Zhang, Lee Graham, Karen Bowker, O. Martin Williams, Alasdair P. MacGowan, Matthew B. Avison

## Abstract

Fluoroquinolone resistance in bacteria is multifactorial, involving target site mutations, reductions in fluoroquinolone entry due to reduced porin production, increased fluoroquinolone efflux, enzymes that modify fluoroquinolones, and Qnr, a DNA mimic that protects the drug target from fluoroquinolone binding. Here we report a comprehensive analysis using transformation and in vitro mutant selection, of the relative importance of each of these mechanisms in fluoroquinolone resistance and non-susceptibility, using *Klebsiella pneumoniae*, one of the most clinically important multi-drug resistant bacterial species known, as a model system. Our improved biological understanding was then used to generate rules that could be predict fluoroquinolone susceptibility in *K. pneumoniae* clinical isolates. Key to the success of this predictive process was the use of liquid chromatography tandem mass spectrometry to measure the abundance of proteins in extracts of cultured bacteria, identifying which sequence variants seen in the whole genome sequence data were functionally important in the context of fluoroquinolone susceptibility.

## INTRODUCTION

The use of whole genome sequencing (WGS) to predict the susceptibility of bacteria to antimicrobials in the clinical setting has recently been subject to a subcommittee report from the European Committee on Antimicrobial Susceptibility Testing (EUCAST).^1^ Aside from issues around universal WGS data availability and quality, particularly with regard to clinical samples with low bacterial titres, the committee considered that a lack of basic biological understanding of antibacterial drug resistance (ABR) was slowing down progress in this area. There have been some successes predicting resistance/susceptibility direct from WGS in key multi-drug resistant Gram negative human pathogens such as *Escherichia coli* and *Klebsiella pneumoniae* when resistance is due to well-known plasmid encoded mechanisms, or when commonly encountered antibiotic target site mutations are present.^2,3^ However, there is a relative paucity of data addressing the issue of how chromosomal mutations, particularly those conferring pleotropic phenotypes affecting envelope permeability, and plasmid encoded resistance mechanisms come together to confer clinically relevant ABR.^1^ In key Gram-negative pathogens, such “multi-factorial” drug resistance is typified by carbapenem resistance where a traditional carbapenem hydrolysing enzyme is not being produced, and fluoroquinolone resistance.^1,4^ The work reported below addresses the latter.

Fluoroquinolones, including norfloxacin, ciprofloxacin, levofloxacin and ofloxacin have a fluorine atom attached to the central quinolone ring system ^5-7^. They have a broad spectrum of activity against both Gram-negative and Gram-positive bacteria and play an important role in clinical applications, particularly for the treatment of urinary tract and respiratory tract infections ^8-10^. The mode of action of fluoroquinolones involves binding to type II topoisomerases - DNA gyrase and topoisomerase IV - to form a drug-enzyme-DNA complex that can break but not re-join DNA during the unwinding of supercoils. This results in lethal double strand breaks in DNA^8,11^.

A single point mutation in the quinolone resistance-determining region (QRDR) of DNA gyrase subunit A (GyrA) is sufficient to confer nalidixic acid resistance in enteric bacteria such as *E. coli* but additional factors are required for fluoroquinolone resistance. For example QRDR mutations in ParC, a subunit of DNA topoisomerase IV and a secondary target of fluoroquinolones ^12^. Fluoroquinolone resistance is also associated with energy dependent efflux due to chromosomal mutations causing overproduction of efflux pumps ^13,14^. The Resistance Nodulation Division (RND) family efflux pump, AcrAB, is frequently seen overproduced in fluoroquinolone resistant clinical Enterobacteriaceae whether caused by mutation in the local repressor, AcrR or through overproduction of an AraC-type transcriptional activator e.g. MarA, or RamA, following mutations in their repressors, MarR or RamR ^15,16^. OqxAB, another RND family multidrug efflux pump, is chromosomally encoded in *K. pneumoniae* but when found in *E. coli* on a conjugative plasmid, pOLA52, it gives resistance to fluoroquinolones and chloramphenicol ^17,18^. OqxAB overproduction in *K. pneumoniae* isolates is associated with mutations in the local repressor OqxR^19^.

Apart from plasmid encoded efflux pumps, the first mobile quinolone resistance determinant was reported on the plasmid pMG252 from a multi-resistant *K. pneumoniae* isolate ^20^. This plasmid-mediated quinolone resistance (PMQR) gene, later named *qnrA1*, encodes a pentapeptide repeat protein which protects DNA gyrase from inhibition by quinolones. Qnr has emerged quickly over the past ten years with other *qnr* gene families such as *qnrB ^21^, qnrC ^22^, qnrS ^23^, qnrD* ^24^ being reported, encoding proteins sharing 43%, 64%, 59%, 48% amino acid identity with QnrA1, respectively, but with a seemingly indistinguishable phenotypic effect. Another common type of PMQR is a variant aminoglycoside acetyltransferase, Aac(6′)-lb-cr, capable of acetylating fluoroquinolones at the amino nitrogen if they have a piperazinyl substituent, e.g. ciprofloxacin^25^.

Whilst it is generally accepted that fluoroquinolone resistance is multi-factorial in Enterobacteriaceae isolates, little work has been done to determine the relative contributions of different mechanisms, individually, and in all possible combinations, to clinically relevant fluoroquinolone susceptibility. One benefit of performing such an analysis is to improve our ability to predict fluoroquinolone susceptibility in Enterobacteriaceae directly from WGS data. Our aim in performing the work set out below was to complete the first extensive analysis of fluoroquinolone susceptibility/resistance in the Enterobacteriaceae using sequential mutant selection, transformation and complementation analysis, with LC-MS/MS proteomics being used to monitor protein production and DNA sequencing being used to identify mutations. We chose to do this using *K. pneumoniae*, because of its growing reputation as the most significant multi-drug resistant Enterobacteriaceae species in human infections. Our aim was to generate a series of “rules” that might be applied to WGS data to predict fluoroquinolone susceptibility in *K. pneumoniae.* We then tested our rules against 40 *K. pneumoniae* clinical isolates.

## MATERIALS AND METHODS

### Bacterial strains and antibiotic susceptibility testing

The isogenic pair *K. pneumoniae* Ecl8 ^26^, Ecl8Δ*ramR* ^27^ and *E. coli* XLIO-Gold (Stratagene) were used throughout. Forty human bloodstream *K. pneumoniae* isolates were studied. Disc susceptibility testing was performed according to CLSI methodology^28^ and interpreted using CLSI performance standards^29^.

### Selection of in vitro K. pneumoniae mutants conferring reduced fluoroquinolone susceptibility and fluoroguinolone resistance

*K. pneumoniae* mutants with reduced fluoroquinolone susceptibility (but perhaps not fluoroquinolone resistance) were generated by plating 100 μl of an overnight culture of wildtype Ecl8 or Ecl8Δ*ramR* on LB agar containing ciprofloxacin or nalidixic acid at lower than the CLSI defined resistance breakpoint concentration. First step mutants were then re-selected using increasing concentrations of ciprofloxacin to generate fluoroquinolone resistant mutants that grew at greater than resistance breakpoint concentrations. Mutants selected were subjected to CLSI disc susceptibility testing. PCR-sequencing of potential resistance causing mutations was performed using primers listed in Error! Reference source not found.

### Cloning genes, transformation and complementation studies of K. pneumoniae Ecl8

The wild-type *oqxR* and *ramR* gene of *K. pneumoniae* Ecl8 were amplified by PCR using the primers set out in **Table S1**. The PCR amplicon was TA cloned into pCR2.1-TOPO (Invitrogen) according to the manufacturer’s instructions. These pCR2.1 inserts were removed using EcoRI (for *oqxR)*, HindIII-HF and XbaI (for *ramR)* and sub-cloned into pK18. The PMQR genes *qnrAl* and *aac(6’)-Ib-cr*, were synthesised (Eurofins Genomics) as defined in **Table S1** to include native promoters and ligated into the pEX-K4 cloning vector. Recombinant plasmids were used to transform *K. pneumoniae* to kanamycin (30 mg/L) resistance using electroporation as standard for laboratory-strain *E. coli.*

### Quantitative analysis of envelope and whole cell proteome via Orbitrap LC-MS/MS

Each bloodstream isolate or transformant was cultured in 50 ml Cation Adjusted Muller-Hinton Broth (Sigma) with appropriate antibiotic selection. Cultures were incubated with shaking (160 rpm) at 37°C until OD_600_ reached 0.5-0.7. Cells in cultures were subjected to total envelope proteomics as described previously.^30^ For total cell proteomics of *K. pneumoniae* bloodstream isolates, cells were cultured in nutrient broth (Oxoid) as above and lysed by sonication; a cycle of 1 sec on, 0.5 sec off for 3 min at amplitude of 63% using a Sonics Vibracell VC-505TM (Sonics and Materials Inc., Newton, Connecticut, USA). Following centrifugation (15 min, 4°C, 8,000 rpm, Sorval RC5B with SS34 rotor), 1 μg of total protein was separated by SDS-PAGE. Gels were run at 200 V until the dye front had moved approximately 1 cm into the separating gel. Proteins in gels were stained with Instant Blue (Expedeon) for 20 min and de-stained in water. The one centimetre of gel lane containing was cut out and proteins subjected to in-gel tryptic digestion using a DigestPro automated digestion unit (Intavis Ltd).

The resulting peptides were fractionated using an Ultimate 3000 nanoHPLC system in line with an LTQ-Orbitrap Velos mass spectrometer (Thermo Scientific).^30^ The raw data files were processed and quantified using Proteome Discoverer software v1.4 (Thermo Scientific) and searched against the UniProt *K. pneumoniae* strain ATCC 700721 / MGH 78578 database (5126 protein entries; UniProt accession 272620) using the SEQUEST (Ver. 28 Rev. 13) algorithm. Protein Area measurements were calculated from peptide peak areas using the Top 3 method ^31^ and were then used to calculate the relative abundance of each protein. Proteins with fewer than three peptide hits were excluded from the analysis. Proteomic analysis was repeated three times, each using a separate batch of cells, except for the clinical isolates, which were run only once.

Data analysis for envelope proteome was as follows: raw protein abundance data were uploaded into Microsoft Excel. A paired T-test was used to calculate the significance of any difference in protein abundance data in pooled data from the two test conditions; where protein abundance was below the level of detection the value was excluded and not set to zero. A p-value of <0.05 was considered significant. The fold change in abundance for each protein in two test conditions was calculated using the averages of absolute abundances for the three biological replicates for the two test conditions. For the clinical isolates where total protein samples were run once each, raw protein abundance for each protein was divided by the average abundance of a ribosomal protein to normalise for sample to sample loading variability.

### Whole genome sequencing and data analysis

Genomes were sequenced by MicrobesNG (Birmingham, UK) on a HiSeq 2500 instrument (Illumina, San Diego, CA, USA). Reads were trimmed using Trimmomatic ^32^ and assembled into contigs using SPAdes 3.10.1 (http://cab.spbu.ru/software/spades/). STs, the presence of resistance genes, and plasmid replicon types were determined using MLST 1.8, ResFinder 2.1,^3^ and PlasmidFinder^33^ on the Center for Genomic Research platform (https://cge.cbs.dtu.dk/services/). Assembled contigs were mapped to reference genome *K. pneumoniae* Ecl8 (GCA_000315385.1) obtained from GenBank using progressive Mauve alignment software.^34^

## RESULTS & DISCUSSION

### Fluoroquinolone resistance in K. pneumoniae occurs in a step-wise manner, requiring combinations of different resistance mechanisms

A total of 192 mutants with reduced fluoroquinolone susceptibility were generated from *K. pneumoniae* Ecl8 and Ecl8Δ*ramR* using nalidixic acid and ciprofloxacin at various concentrations. Characterisation of these mutants revealed that selection of *in vitro* fluoroquinolone resistance occurs in a step-wise manner with the accumulation of different mutations at each step.

Among the 29 representative mutants that were extensively characterised, 16 had quinolone target site mutations in *gyrA* clustered into four different alleles, each encoding a single amino acid substitution within the QRDR: Ser83Phe, Ser83Tyr, Asp87Tyr or Gly81Cys. Multiple mutations in *gyrA* or mutations in other topoisomerase genes, e.g. *parC* were not identified. Mutations in known regulators of drug efflux were also seen. OqxR mutations were found in 12/27 fully characterised mutants, including frameshift mutations and nonsense and missense mutations at various positions **(Table S2)**. In contrast, only a single (Thr124Pro) RamR mutation was found, and no mutations in AcrR were identified. The roles of two PMQR genes were also assessed: *qnr* (represented by the *qnrA1* variant) and *aac(6’)-lb-cr*, both being provided on a vector with expression driven by their native promoters.

It was observed that whilst they all reduced fluoroquinolone susceptibility, single acquisition events; whether GyrA mutations, RamR or OqxR mutations, or carriage of a PMQR gene did not confer clinically relevant fluoroquinolone non-susceptibility (i.e. defined as intermediate resistance or resistance to ciprofloxacin, by reference to the relevant breakpoints) in *K. pneumoniae* Ecl8. The largest reduction in susceptibility seen was in *oqxR* loss of function mutants (Error! Reference source not found.).

Combinations of an *oqxR* or *ramR* plus a *gyrA* mutation or of an *oqxR* or *ramR* mutation plus carriage of *qnr* (Error! Reference source not found.) were the only double combinations that conferred clinically relevant ciprofloxacin non-susceptibility in Ecl8. In 10 triple and five quadruple combinations tested, all conferred fluoroquinolone non-susceptibility in Ecl8 except the *ramR/oqxR* double mutant plus *aac(6')-Ib-cr* **(Table S5, S6),** which is known to be a weak PMQR determinant and is only capable of acetylating ciprofloxacin and norfloxacin, which have an unsubstituted piperazinyl group.^25^

### Relative impact of OqxR and RamR mutation to efflux pump production

Complementation of the *oqxR* and *ramR* mutations with wild type genes *in trans* confirmed that loss of OqxR has a stronger effect on fluoroquinolone susceptibility than RamR loss **(Table S7)**. To investigate the mechanism behind this difference, we used Orbitrap LC-MS/MS proteomics to define the impact of these mutations on the production of porins and efflux pumps previously implicated in antimicrobial drug resistance in *K. pneumoniae.* The OqxA and OqxB efflux pump proteins were found to be below the detection level in wild-type cells, however, they are detectable in both the *ramR* and *oqxR* mutants with upregulation being almost two orders of magnitude greater in the latter (Error! Reference source not found**.c,** Error! Reference source not found**.d).** By contrast, significant changes in AcrA and AcrB efflux pump protein abundance were not observed in the *oqxR* mutant but AcrAB were overproduced in the *ramR* mutant (Error! Reference source not found**.a,** Error! Reference source not found.**b)** with similar fold increases to those reported recently for the Ecl8 *ramR* deletion mutant.^37^ TolC is likely to work as the outer membrane protein for both OqxAB and AcrAB, and TolC levels were significantly upregulated in both mutants relative to wild type (Error! Reference source not found**.e)**. Downregulation of the OmpK35 (OmpF) porin was equally observed in both mutants: 0.43-fold, p=0.005 *{oqxR* mutant), 0.41-fold p=0.005 *{ramR* mutant) **(Figure 1f)**. Significant changes in abundance of the OmpK36 (OmpC) porin were not observed in either mutant (Error! Reference source not found**.g))**.

**Figure 1:**
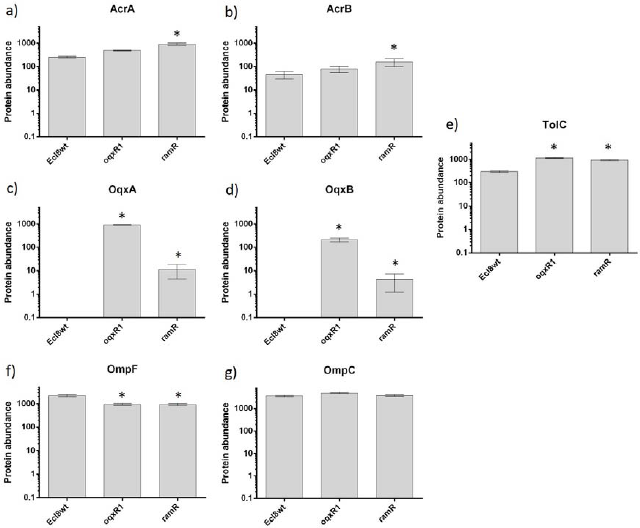
Envelope proteome changes in *oqxR* and *ramR* mutant relative to Ecl8wt for each protein. Values reported in the bar graph are absolute abundance values (x10^−7^). Data, are the mean (n=3); error bars are standard error of the mean. A star (*) above a bar indicates significant changes following: fold change ≥ 2 fold, p<0.05 for a T-test comparing absolute protein abundance data, n=3.

### Predicting fluoroquinolone susceptibility in K. pneumoniae clinical isolates from whole genome sequencing

The results from *in vitro K. pneumoniae* Ecl8 mutant and transformant characterisation (Error! Reference source not found, **to S6)** were used to generate a set of rules to predict fluoroquinolone susceptibility in *K. pneumoniae* as shown in Error! Reference source not found. The rules were then applied to 10 *K. pneumoniae* clinical isolates based on WGS data. The presence of PMQR genes was defined as a binary result from the WGS data, given that all genes were fully intact. The presence of QRDR mutations in GyrA and of SNPs affecting the amino acid sequence of RamR and OqxR could be read directly from the WGS data. To confirm whether the regulatory SNPs activated efflux pump production, or were random genetic drift, whole cell LC-MS/MS proteomics was performed. Clinical isolates carrying Thr141Ile and 194K insertion mutations in RamR (6/10 clinical isolates carrying one or both, suggestive of random genetic drift) did not hyperproduce AcrA and AcrB, as would be expected from a true RamR loss of function mutation **(figure 1),** so these variants were considered functionally wild-type. Two out of 10 isolates carried OqxR mutations, and both were confirmed to hyperproduce OqxB. Interestingly, one of these mutants (KP9) had also entirely lost the *ramR/ramA* locus as part of a ~70 kb deletion in the chromosome. We identified double *gyrA* mutations in 2/10 isolates and *parC* mutations in 4/10 isolates, something not seen in any of the Ecl8 reduced susceptibility/resistant mutants selected in vitro.

Applying our predictive rules **(Table S8),** WGS information for the 10 clinical isolates, with proteomics being used to confirm/deny suspected regulatory lesions, allowed correct prediction of ciprofloxacin susceptibility, as confirmed using disc susceptibility testing, in 7/10 isolates. The three remaining isolates were incorrectly predicted to be ciprofloxacin susceptible as defined by Rule 15, which is carriage of Qnr and the Aac-6'-Ib-cr in an otherwise wild-type background **(Table S8)**. The isolates were ciprofloxacin non-susceptible, but susceptible to all other fluoroquinolones tested **(Table 1)**. We hypothesised that these isolates have reduced envelope permeability compared with Ecl8, the isolate used to define our predictive rules **(Table S8)**. Fluorescent dye accumulation assays **(Fig. 2A, B)** revealed that the shape of the dye accumulation curve was similar to that seen previously upon *micF* overexpression in Ecl8, i.e. reduced OmpF porin production.^35^ This retards entry of the dye, but does not prevent accumulation of the dye over time **(Fig. 2A, B)**. This phenotype is in stark contrast to efflux mediated reduced envelope permeability, which gives a persistent level of reduced dye accumulation, as seen in the Ecl8Δ*ramR* mutant previously ^35^, and in clinical isolate KP21, a clinical isolate with a *ramR* mutation from our collection **(Table 2, Fig. 2C)**. Importantly, the Rule 15 isolates produce Aac-6'-Ib-cr, which modifies ciprofloxacin ^25^. Hence, even a permeability mutation that reduces the initial rate of entry of ciprofloxacin into the cell **(Fig. 2A, B)** can enhance the ability of a ciprofloxacin modifying enzyme to reduce active ciprofloxacin accumulation. We noticed that even isolates defined as genotypically wild-type relative to Ecl8 revealed smaller inhibition zone diameters for fluoroquinolones than Ecl8 **(Table 2; Table S3)**. So, we predicted that, like the Rule 15 isolates, they too have reduced envelope permeability. This proved to be correct and transformation of one of these isolates, KP46 **(Fig. 2D, Table 2)** with a plasmid carrying *qnr* and *aac-6'-Ib-cr* (i.e. reconstituting the Rule 15 genotype, but in a background with reduced permeability), generated a transformant that was ciprofloxacin non-susceptible but susceptible to all other fluoroquinolones tested **(Table S9),** as predicted from the observed phenotype of the clinical isolates genotypically identified as following Rule 15 **(Table 1)**. In order to confirm the suspected porin deficiency in these clinical isolates, we analysed the LC-MS/MS data and calculated an OmpF:OmpC ratio for each isolate. For Ecl8 during growth in the same medium used to perform the proteomic analyses on our clinical isolates (nutrient broth), the OmpF:OmpC ratio is approximately 1:1.^35^ For the 10 initial test clinical isolates, the OmpF:OmpC ratio was in a range of 0.25:1 to 0.40:1, or zero **(Table 1)**. The two isolates that have no detectable OmpF in the LC-MS/MS, KP3 and KP4 have Trp316STOP and Gly208FRAMESHIFT mutations, respectively, according to WGS. This observed reduced OmpF:OmpC ratio in the clinical isolates relative to Ecl8 rationalised the observed envelope permeability assay data **(Fig. 2)** and supported our hypothesis that the rate of ciprofloxacin entry is retarded in Rule 15 isolates relative to Ecl8, creating a ciprofloxacin non-susceptible phenotype **(Table. S9)**.

**Figure 2:**
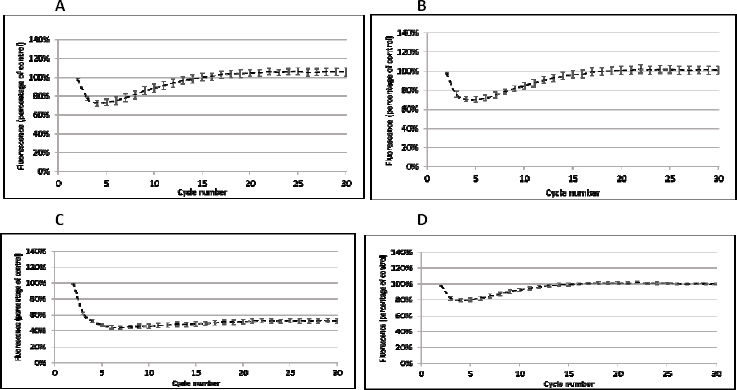
The accumulation of H33342 dye over a 30 cycle (45 minute) incubation period by *K. pneumoniae* clinical isolates: (A) KP7, (B) KP8, (C) KP21, (D) KP46. In each case, permeability was compared with Ecl8 (set to 100%) in cells grown in MHB. Each line shows mean data for three biological replicates with 8 technical replicates in each and error bars define the standard error of the mean (SEM)

**Figure 3:**
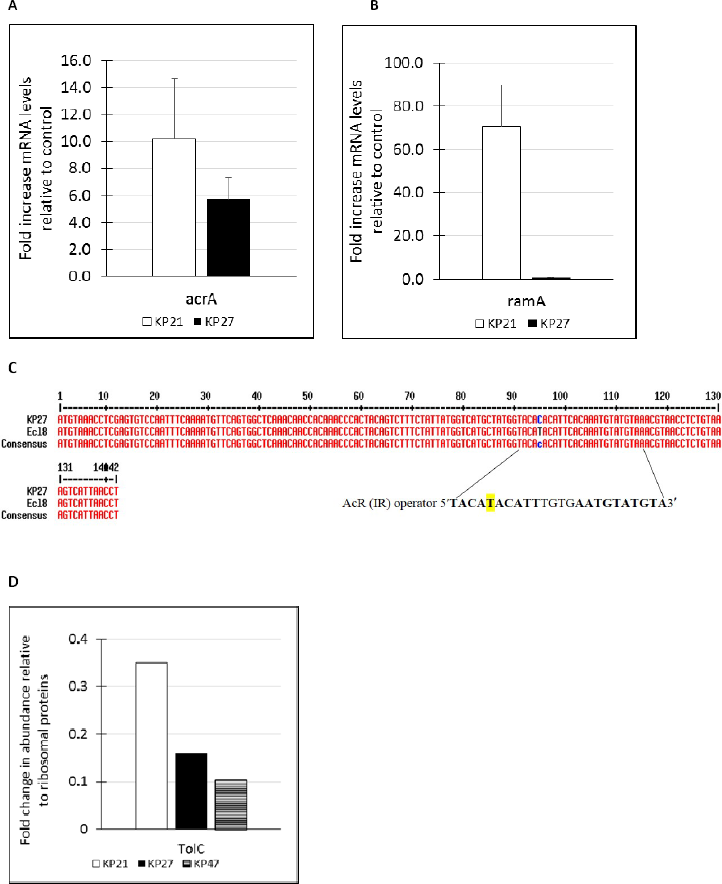
Expression of (A) *arcA* and (B), *ramA* in *K. pneumoniae* clinical isolates KP21 *(ramR* mutant) and KP27 *(ramR* wild-type) both normalised to expression in a control isolate KP17, using qRT-PCR. Data are presented as means +/− SEM, *n*=3 preparations of RNA (C) shows that a single nucleotide polymorphism is present in the putative AcrR binding site upstream of acrAB in KP27. (D) shows TolC protein abundance in KP21, KP27 and the control isolate KP47, data being normalised in each sample to the average ribosomal protein.

**Table 1.** Combination of disc test, proteomics, WGS and predictions of fluoroquinolone susceptibility for *K. pneumoniae* clinical isolates

| Isolate | GyrA | OqxR | OqxB | RamR | AcrA, B | *qnr* | *aac(6')-Ib-cr* | Prediction | CIP<br>(5) | LEV<br>(5) | NOR<br>(10) | OFL<br>(5) | ParC | OmpF:<br>OmpC<br>ratio | New Prediction |
| --- | --- | --- | --- | --- | --- | --- | --- | --- | --- | --- | --- | --- | --- | --- | --- |
|  | Mutation | Mutation | Abundance | Mutation | Abundance | Presence | Presence | (Rule) | Zone | Zone | Zone | Zone | Mutation |  | (Rule) |
| KP1 | - | - | ND | - | 0.07, 0.04 | S | - | S (4) | 24 | 21 | 23 | 19 | - | 0.30 | S (4) |
| KP2 | - | - | ND | *Thr141Ile | 0.10, 0.03 | B, S | + | S (15) | 18 | 22 | 20 | 19 | - | 0.28 | NS (15*) |
| KP3 | Ser83Ile | - | ND | *Thr141Ile | 0.11, 0.03 | B | + | NS (21) | 6 | 10 | 6 | 7 | Ser80Ile | 0.00 | NS (21) |
| KP4 | Ser83Tyr<br>Asp87Gly | - | ND | Lys9Ile<br>*Thr141Ile | 0.26, 0.2 | S | - | NS (19) | 6 | 6 | 6 | 6 | Ser80Ile | 0.00 | NS (19) |
| KP5 | - | *oqxR*<br>deleted | 0.23 | *Thr141Ile<br>*194K | 0.08, 0.07 | - | - | S (2) | 22 | 20 | 19 | 17 | - | 0.28 | S (2) |
| KP6 | Ser83Tyr | - | ND | - | 0.10, 0.03 | B | + | NS (21) | 13 | 18 | 13 | 15 | - | 0.25 | NS (21) |
| KP7 | - | - | ND | - | 0.13, 0.06 | B | + | S (15) | 16 | 21 | 19 | 18 | - | 0.25 | NS (15*) |
| KP8 | - | - | ND | *Thr141Ile | 0.13, 0.04 | B | + | S (15) | 18 | 22 | 19 | 19 | - | 0.25 | NS (15*) |
| KP9 | Ser83Phe<br>Asp87Ala | Glu24STOP | 0.06 | *ramR/ramA*<br>deleted | 0.08, 0.03 | - | - | NS (6) | 10 | 14 | 8 | 11 | Ser80Ile | 0.40 | NS (6) |
| KP10 | Ser83Ile | - | ND | *Thr141Ile | 0.10, 0.02 | B | + | NS (21) | 6 | 11 | 6 | 9 | Ser80Ile | 0.26 | NS (21) |

Values reported for zone diameter are the means of three repetitions rounded to the nearest integer
Zones shaded bright blue are resistant, pale blue are intermediate resistant (non-susceptible) according to susceptibility breakpoints set by the CLSI
Zones shaded green and red showed success and failure in predicting FQ susceptibility/resistance respectively
\*Changes in amino acid due to genetic variation. Unlikely to have any effect on gene function
+, gene is present, -, mutations not detected or gene is absent, ND; protein abundance not detected

**Table 2.** Combination of disc test, proteomics, genome sequencing and predictions of fluoroquinolone susceptibility for *K. pneumoniae* clinical isolates

| Isolate | GyrA<br>Mutation | ParC<br>Mutation | OqxR<br>Mutation | OqxB<br>Abundance | RamR<br>Mutation | AcrA, B<br>Abundance | *qnr*<br>Presence | *aac(6')-Ib-cr*<br>Presence | OmpF:OmpC<br>ratio | Prediction<br>(Rule) | CIP<br>(5)<br>Zone | LEV<br>(5)<br>Zone | NOR<br>(10)<br>Zone | OFL<br>(5)<br>Zone |
| --- | --- | --- | --- | --- | --- | --- | --- | --- | --- | --- | --- | --- | --- | --- |
| KP11 | Ser83Ile | Ser80Ile | - | ND | *Thr141Ile<br>Met184Val<br>*194K | 0.24, 0.09 | S | - | 0.00 | NS (19) | 6 | 6 | 6 | 6 |
| KP12 | Ser83Ile | Ser80Ile | Asp3Tyr | 0.11 | *Thr141Ile | 0.14, 0.04 | - | - | 0.00 | NS (6) | 6 | 6 | 6 | 6 |
| KP13 | Ser83Ile | Ser80Ile | - | ND | Lys63FS | 0.39, 0.2 | B, S | + | 0.00 | NS (29) | 6 | 6 | 6 | 6 |
| KP14 | - | - | - | ND | *ramR/ramA*<br>deleted | 0.09, 0.04 | B | + | 0.36 | NS (15*) | 20 | 24 | 22 | 22 |
| KP15 | - | - | - | ND | - | 0.14, 0.03 | S | + | 0.21 | NS (15*) | 19 | 23 | 19 | 21 |
| KP16 | Ser83Ile | Ser80Ile | - | ND | *Thr141Ile | 0.11, 0.03 | S | - | 0.40 | S (8) | 9 | 9 | 10 | 6 |
| KP17 | Ser83Ile | Ser80Ile | - | ND | *Thr141Ile | 0.11, 0.04 | - | - | 0.35 | S (1) | 17 | 17 | 15 | 14 |
| KP18 | - | - | - | ND | Ala19Val<br>*Thr141Ile | 0.12, 0.05 | S | + | 0.36 | NS (15*) | 18 | 22 | 17 | 19 |
| KP19 | - | - | - | ND | *Thr141Ile<br>*194K | 0.1, 0.03 | B | + | 0.42 | NS (15*) | 19 | 23 | 20 | 21 |
| KP20 | - | - | Ala19Val | 0.08 | *Thr141Ile | 0.11, 0.04 | B | - | 0.00 | NS (11) | 14 | 13 | 14 | 10 |
| KP21 | - | - | - | ND | Arg44FS | 0.41, 0.23 | S | - | 0.00 | NS (13) | 14 | 13 | 13 | 10 |
| KP22 | - | - | - | ND | *Thr141Ile<br>*194K | 0.09, 0.06 | B | + | 0.45 | NS (15*) | 19 | 23 | 20 | 20 |
| KP23 | - | - | - | ND | *Thr141Ile<br>*194K | 0.05, 0.03 | B | + | 0.41 | NS (15*) | 19 | 23 | 20 | 20 |
| KP24 | - | - | - | ND | - | 0.13, 0.07 | B | + | 0.34 | NS (15*) | 19 | 23 | 20 | 20 |
| KP25 | - | - | - | ND | *Thr141Ile | 0.12, 0.09 | B | + | 0.29 | NS (15*) | 19 | 23 | 20 | 20 |
| KP26 | - | - | - | ND | *Thr141Ile | 0.08, 0.02 | B | + | 0.33 | NS (15*) | 19 | 23 | 20 | 20 |
| KP27 | Ser83Tyr<br>Asp87Phe | Ser80Ile | - | ND | *Thr141Ile | 0.36, 0.15 | - | + | 0.00 | NS (9) | 6 | 11 | 6 | 8 |
| KP28 | - | - | Leu76FS | 0.13 | *Thr141Ile | 0.12, 0.04 | S | - | 0.00 | NS (11) | 13 | 12 | 11 | 8 |
| KP30 | Ser83Ile | Ser80Ile | Val130Ala | 0.14 | Ala40Val<br>*Thr141Ile<br>*194K | 0.87, 0.33 | - | - | 0.00 | NS (16) | 6 | 6 | 6 | 6 |
| KP31 | - | - | - | ND | *Thr141Ile | 0.08, 0.04 | - | - | 0.43 | S (WT) | 31 | 29 | 28 | 27 |
| KP32 | - | - | - | ND | - | 0.08, 0.04 | - | - | 0.40 | S (WT) | 32 | 28 | 30 | 26 |
| KP33 | - | - | - | ND | *Thr141Ile | 0.08, 0.03 | - | - | 0.37 | S (WT) | 34 | 30 | 31 | 28 |
| PK34 | - | - | - | ND | *Thr141Ile | 0.16, 0.07 | - | + | 0.36 | S (5) | 26 | 28 | 24 | 26 |
| KP38 | - | - | - | ND | *Thr141Ile | 0.05, 0.31 | - | - | 0.00 | S (WT) | 31 | 29 | 28 | 27 |
| KP40 | - | - | - | ND | Ala19Val<br>*Thr141Ile | 0.03, 0.03 | - | - | 0.36 | S (WT) | 32 | 29 | 31 | 27 |
| KP46 | - | - | - | ND | - | 0.03, 0.01 | - | - | 0.34 | S (WT) | 34 | 30 | 30 | 27 |
| KP47 | - | - | - | ND | - | 0.10, 0.04 | - | - | 0.32 | S (WT) | 33 | 29 | 31 | 27 |
| KP48 | - | - | - | ND | - | 0.08, 0.04 | - | - | 0.35 | S (WT) | 34 | 30 | 30 | 28 |
| KP50 | - | - | - | ND | Ile106Ser<br>*Thr141Ile | 0.05, 0.03 | - | - | 0.45 | S (WT) | 34 | 31 | 30 | 30 |
| KP59 | - | - | - | ND | Thr50Ala | 0.21, 0.11 | - | - | 0.41 | S (3) | 33 | 28 | 31 | 26 |

Values reported for zone diameter are the means of three repetitions rounded to the nearest integer
Zones shaded bright blue are resistant, pale blue are intermediate resistant (non-susceptible) according to susceptibility breakpoints set by the CLSI
Zones shaded green and red showed success and failure in predicting FQ susceptibility/resistance respectively
\*Changes in amino acid due to genetic variation. Unlikely to have any effect on gene function
+, gene is present, -, mutations not detected or gene is absent, ND; protein abundance not detected

Based on the findings concerning Rule 15, we revised our rules to include Rule 15* which predicts ciprofloxacin non-susceptibility, but only applies if the OmpF:OmpC ratio is ≤0.5. We then tested these adapted rules versus 30 additional clinical isolates. In this case, we were successful in predicting ciprofloxacin susceptibility/non-susceptibility in 28/30 isolates, with all 30 isolates showing reduced OmpF:OmpC ratio relative to Ecl8 and Rule 15 isolates being the most common **(Table 2)**. The two isolates we incorrectly predicted to be ciprofloxacin susceptible each carried more than one Gyrase/Topoisomerase mutation, never seen in our Ecl8 mutants but relatively common in our clinical isolates. We would presume therefore that these double mutations are the reason for ciprofloxacin non-susceptibility, but it is important to note that they also have a reduced OmpF:OmpC ratio.

Interestingly isolate KP27 has a wild-type *ramR* sequence, but the proteomics showed hyper-production of AcrAB **(Table 2)**. This was confirmed using qRT-PCR for *acrA* relative to a wild-type control isolate, KP47, but *ramA* was not over-expressed in KP27, validating the observation that RamR is wild-type **(Fig. S3A, B, Table 2)**. WGS identified a mutation in the *acrR-acrA* intergenic region in KP27, causing a T-C transition at the 5^th^ position of the first AcrR binding motif (5’ TACATACATT3’) **(Fig. S3C),** predicted to derepresses *acrAB* expression.^36^ Interestingly, according to proteomics, KP27 does not overproduce TolC, relative to wild-type clinical isolate KP47. This contrasts with isolate KP21, for example, which is a *ramR* loss of function mutant **(Fig. S3A, B, D, Table 2)**. This difference is as expected, because *acrAB* and *tolC* are unlinked and the latter is not controlled by AcrR, but both are controlled by RamA.^37^

## CONCLUSIONS

We have demonstrated for the first time in *K. pneumoniae*, step-wise combination of resistance mechanisms as well as characterising the importance of each mechanism and the interplay between them in conferring fluoroquinolone resistance. Our analysis showed mutations in GyrA QRDR, efflux pump overproduction and presence of PMQR on their own do not confer clinically relevant fluoroquinolone resistance. Combinations of OqxR or RamR mutation conferring OqxAB-TolC or AcrAB-TolC overproduction, respectively, together with either GyrA mutation or Qnr PMQR were the only double combination of two mechanisms conferring fluoroquinolone resistance. However, we did not see any multiple gyrase/topoisomerase mutants, so their combined effect on fluoroquinolone susceptibility cannot be definitively confirmed. It was interesting to find this, since we screened 192 individual mutants, but loss of function mutations in efflux pump regulators are higher frequency events (10^3^-10^4^ times more likely) than specific target site mutations, so perhaps this is not a surprise. It was surprising, however, to find 11/40 gyrase/topoisomerase multiple mutants in the clinical isolates and 11/40 efflux pump regulatory mutants. Perhaps, there is a greater fitness cost associated with efflux pump overproducing mutations than target site mutations, so in the real world, target site mutations are relatively more common than their mutation frequency might lead us to expect.

Using this information, we have successfully applied WGS to predict ciprofloxacin resistance in *K. pneumoniae* bloodstream isolates. LC-MS/MS proteomics was the key to understanding the relative importance of different SNPs seen in the efflux pump regulators, and the more phenotypically relevant SNPs we can identify, the more complex rules for predicting fluoroquinolone resistance directly from WGS can be developed. Because our initial rules for predicting resistance were defined using Ecl8, a domesticated strain, we incorrectly predicted ciprofloxacin susceptibility in isolates carrying both PMQRs studied, but no mutation in efflux pump regulators or gyrase/topoisomerase genes. Indeed, the clinical isolates have reduced envelope permeability relative to Ecl8, due to a downregulation of OmpF, and a consequent reduction in the OmpF:OmpC porin ratio. This is another factor for which proteomics enhanced the predictive power of WGS. Once we understand the genetic basis for this reduced OmpF:OmpC ratio we can factor it into our predictive rules, and will be able to confirm whether this genetic difference is essential for multiple gyrase/topoisomerase mutations to confer non-susceptibility to fluoroquinolones.

Overall, this work as enhanced our biological understanding of fluoroquinolone resistance in *K. pneumoniae* and will improve our ability to predict susceptibility in clinical isolates based on WGS data. Particularly, it shows the value of using proteomics to clarify genotype to phenotype relationships, and to bridge the gap between WGS and antimicrobial susceptibility testing where mechanisms of resistance are multi-factorial and complex.

## Acknowledgements

Genome sequencing was provided by MicrobesNG (http://www.microbesng.uk), which is supported by the BBSRC (grant number BB/L024209/1).

## Transparency Declaration

None to declare - All authors.

